# Corticomuscular control of walking in older people and people with Parkinson’s disease

**DOI:** 10.1101/628016

**Authors:** Luisa Roeder, Tjeerd W Boonstra, Graham K Kerr

## Abstract

Changes in human gait that result from ageing or neurodegenerative diseases are multifactorial. Here we assess the effects of age and Parkinson’s disease (PD) on corticospinal control in electrophysiological activity recorded during treadmill and overground walking. Electroencephalography (EEG) from 10 electrodes and electromyography (EMG) from two leg muscles were acquired from 22 healthy young, 24 healthy older and 20 adults with PD. Event-related power, corticomuscular coherence (CMC) and inter-trial coherence were assessed for EEG from bilateral sensorimotor cortices and EMG from tibialis anterior muscles during the double support phase of the gait cycle. CMC and EMG power in the low beta band (13-21 Hz) was significantly decreased in older and PD participants compared to young people, but there was no difference between older and PD groups. Older and PD participants spent shorter time in the swing phase than young individuals. These findings indicate age-related changes in the temporal coordination of gait. The decrease in beta CMC suggests reduced cortical input to spinal motor neurons in older people during the double support phase. We also observed multiple changes in electrophysiological measures at high beta and low gamma frequencies during treadmill compared to overground walking, indicating task-dependent differences in corticospinal locomotor control.

## Introduction

Human gait is known to change with age and to be affected by neurodegenerative diseases such as Parkinson’s disease (PD). Changes in gait often affect mobility and activities of daily living, and increase the risk for falls, disability, morbidity, and mortality in these populations^1–3^. Mechanical and dynamical changes in gait have been well documented. Older adults are known to walk slower, take shorter strides, spend more time in stance and double support, use their hip extensors more, and their ankle plantar flexors and knee extensors less than young individuals^4–10^. Movement kinematics, including joint moments and powers at the ankle, and ground reaction forces, also change with ageing^11^. PD gait is characterised by slowness, increased variability and impaired postural control^1,12–16^. Many stride parameters are altered in people with PD compared to age-matched controls, including decreased walking velocity, stride length, cadence, arm swing, head-trunk control and single support time^17,18^. Spatial and temporal gait parameters are also more variable^15^ and more asymmetric^1^. In addition, PD patients may experience episodic gait disturbances such as freezing or festination of gait^1,12,19^. In recent years, treadmill training has been increasingly used to improve gait in PD and its therapeutic effects are greatly discussed. Beneficial effects of treadmill training have been reported for gait speed, step length, stride variability, balance and freezing^20–25^. Notably, treadmill walking has been shown to alter gait kinematics and leg muscle activation compared to overground walking in both healthy adults^26–29^ and in people with PD^30–34^.

The neural mechanisms associated with age- and disease-related gait changes have been investigated using neuroimaging techniques, such as functional magnetic resonance imaging (fMRI), near-infrared spectroscopy (NIRS) and electroencephalography (EEG). Studies using these imaging techniques have shown that gait-related brain activity differs in older adults and people with PD^35–40^. For example, older adults exhibit greater activity in various cortical regions including motor, somatosensory, visual and frontal cortices than young individuals during gait imagery as revealed by fMRI^41–43^. In people with PD, fMRI studies have shown that neural activity during gait imagery tasks is altered in various brain regions compared to age-matched controls^44–46^, and that freezing of gait is associated with impaired functional connectivity between neural networks^47,48^. Moreover, regional cerebral blood flow in the supplementary motor area (SMA) was found to be decreased during treadmill walking in people with PD compared to healthy controls as detected by SPECT^49^. While these studies indicate changes in gait-related brain activity that occur with ageing and PD, they are limited to imagined gait (or recordings after the performance of walking tasks) and may not apply to real walking as participants cannot move during a fMRI scan. NIRS, on the other hand, is a mobile imaging technique that allows capturing changes in oxygenated haemoglobin levels during walking. Using NIRS, Harada et al.^50^ found that the SMA, sensorimotor cortices and the prefrontal cortex are involved in gait control in healthy older people. Prefrontal cortical activity has been observed to decrease during walking with additional cognitive tasks in older but not in young people^51^ and has been observed to predict falls^52^. In people with PD, oxygenated haemoglobin levels over frontal and pre-frontal regions have been reported to be sensitive to freezing of gait (FOG) episodes, walking with obstacle crossings and dual-task walking^53–56^.

EEG can be used to record electrical activity of the brain during walking with high temporal resolution. EEG signals can be analysed in the time (event-related potentials) and frequency (oscillatory activity) domain to investigate cortical control of gait. EEG acquired during treadmill walking has revealed delayed and diminished event-related potentials in response to a cognitive go/no-go task in older compared to young adults^57^. In people with PD, movement-related cortical potentials during a gait initiation task were also reduced compared to healthy young people^58^. Time-frequency analysis revealed that cortical oscillations were modulated by age and PD. EEG gamma power over frontal cortices was enhanced in older people during dual-task walking^59^. Theta and low-gamma power was increased during turning with freezing of gait in people with PD compared to normal fluent turning in a number of cortical regions^60,61^. Interestingly, previous research has also demonstrated that people with PD show abnormalities in beta oscillatory activity in the basal ganglia (subthalamic nucleus, STN) and cerebral cortex at rest as well as during movement^62–67^. For instance, at rest in an untreated state (after a medication wash-out period and STN deep brain stimulating (DBS) electrodes switched off), beta power in the STN and beta activity amongst basal ganglia-cortical networks was increased compared to an optimally medicated state and STN DBS switched on^68–72^. Notably, walking and cycling was found to decrease STN beta power in people with PD^73–75^. Moreover, stepping movements have been shown to modulate STN beta power relative to the movement phase (suppressed during contralateral foot lift)^74^. Some disturbances in gait-related brain activity may be due to the dopamine deficit in the basal ganglia–thalamo–cortical circuit in PD, but other pathways, including cortical cholinergic deficits, may also contribute to PD gait disorders^1,37,76^.

Although both the changes in brain activity and movement kinematics during gait in older people and PD have been extensively documented, it is not clearly understood how these changes are related. How do the changes in brain activity result in the specific gait changes observed with ageing and in PD? Investigating functional interactions between the central and peripheral nervous system may help to address this research question. For example, corticomuscular (EEG-EMG) coherence has revealed correlated activity in the beta band that likely reflects efferent activity from corticospinal projections^77,78^. While corticomuscular coherence has been mainly investigated during precision grip tasks, it has also been assessed during walking^79^. We recently showed that in young healthy adults corticomuscular coherence is observed during the double support phase of the gait cycle and largely absent during the swing phase, and that treadmill walking reduces corticomuscular coherence at high beta frequencies compared to walking overground^80^. Using directed connectivity analysis, Artoni et al.^81^ showed that the functional interactions during normal walking primarily reflect unidirectional drive from contralateral motor cortex to muscles. Changes in corticomuscular coherence with ageing and PD has not yet been investigated during walking, but a few studies have investigated it in other tasks. During cyclical ankle movements – as a simulation of rhythmic natural walking – corticomuscular coherence was reduced in older compared to young participants^82^, but it was not different between people with PD and age-matched controls even though movement performance varied between these groups^83^. Notably, beta-band coherence was enhanced in older compared to young participants during abduction of the index finger^84^, although others reported it reduced during elbow flexion^85,86^. Interestingly, levels of corticomuscular beta-band coherence during forearm contractions were found to be similar in people with PD and age-matched controls^87,88^. Yet, STN DBS has been observed to enhance corticomuscular beta-band coherence in people with PD during a precision grip task^89^, and levodopa medication during forearm contractions has been found to reduce it^87^.

In this study we aimed to investigate whether corticomuscular coherence during walking is affected in older people and people with PD. We therefore collected ambulatory EEG and EMG during overground and treadmill walking in three different groups: young, old and PD. We assessed corticomuscular coherence during the gait cycle in both gait modalities as well as other spectral measures that have previously been investigated during gait^59,80, 90–93^. If the changes in gait observed with ageing and/or PD reflect, at least in part, deficient cortical control due to neurodegeneration in the brain, we expect that corticomuscular coherence will be affected in the older and PD groups and may be differently modulated by overground and treadmill gait. By investigating functional interactions between brain and muscle we aim to contribute to a better understanding of the changes in cortical control of gait with ageing and PD.

## Results

We investigated corticomuscular activity and coupling in 24 healthy young adults, 24 healthy older adults and 21 individuals with PD, while participants walked at their preferred speed and EEG from 10 cortical sites and EMG from the tibialis anterior (TA) muscles were recorded. In particular, we compared changes in spectral power, inter-trial coherence (ITC) and corticomuscular coherence (CMC) at two bipolar derivatives over bilateral sensorimotor cortices between groups to investigate changes in the neural control of gait with ageing and PD. In addition, temporal gait parameters were calculated from footswitch recordings.

In the young and PD groups, data of some participants were excluded from the spectral analysis (two from healthy young group, one from PD overground, three from PD treadmill) due to either excessive EEG artefacts across all channels and conditions (large-amplitude movement artefacts (>300µV) occurring with regular rhythmicity in each gait cycle throughout the entire record), or due to problems with the footswitch recordings (fewer than 100 heel strike triggers). Hence, spectral measures, demographics, clinical and gait assessments were based on n = 22 for the younger group, n = 24 for the older group, and n = 20 for PD group (n = 18 for spectral measures during treadmill walking). In addition, some data were missing for particular outcome measures: in the older group data of one participant were missing for body weight, visual function, and tactile sensitivity (n = 23 for these measures); in the PD group data of one participant were missing for the Bailey Lovie – low contrast sensitivity vision test (n = 19 for this measure).

### Group characteristics

The mean (SD) age of healthy young participants was 25.9 (3.2) years, they had a body height of 170.4 (9.5) cm, and body weight of 68.8 (12.1) kg; 50% were male. Participants in the old and PD groups were of similar age (old: 65.1 (7.8) years, PD: 67.4 (7.3) years), height (old: 169.2 (8.2) cm, PD: 170.3 (9.2) cm), weight (old: 75.2 (14.2) kg, PD: 74.9 (14.8) kg) and gender balance (50% male in the older and 60% in the PD group). PD participants were in the early stages of the disease (Hoehn & Yahr score 1.3) and had an average MDS-UPDRS score of 38.8, indicating a mild to moderate disease severity. The results of the Gait & Falls Questionnaire (GFQ) demonstrated that the PD participants experience only mild gait impairments (Table 1).

**Table 1.**
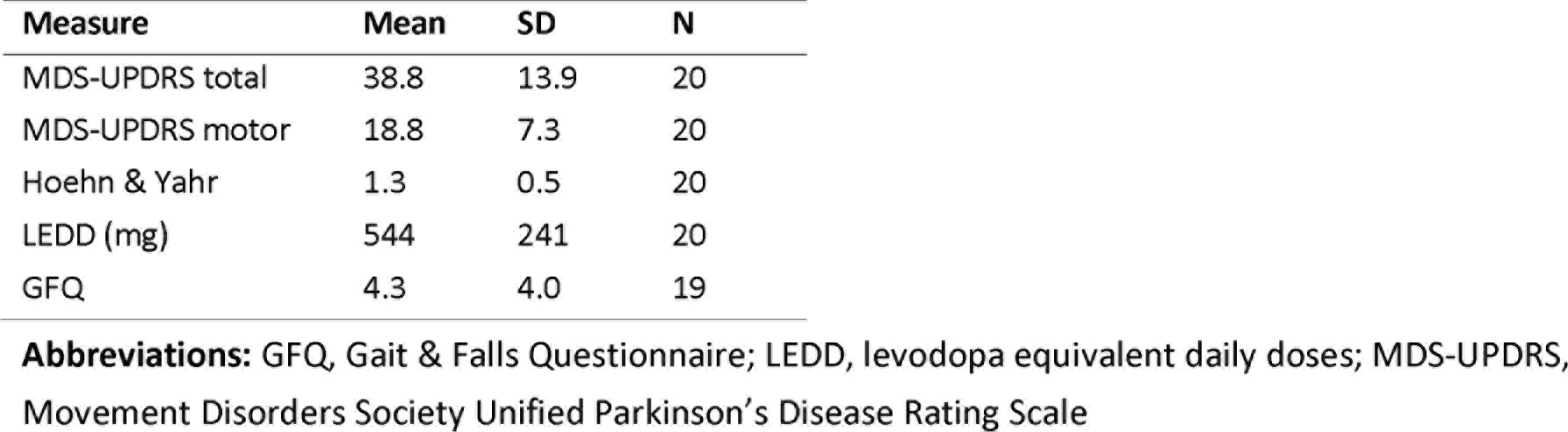
Clinical characteristics of PD group

The PD and healthy older control group had similar levels of cognitive function, no cognitive decline, and all were well above recognised cut-offs for dementia^94^. Both groups had similar levels of tactile sensitivity on the soles of their feet (Table 2). Tactile sensitivity was measured using a Semmes-Weinstein pressure aesthesiometer, which contained eight nylon monofilaments corresponding to eight thresholds^95^. High and low contrast visual acuity as well as contrast sensitivity (Pelli Robson; Melbourne Edge Test) were impaired in people with PD compared to controls (p ≤ 0.03). The Activities-specific Balance Confidence (ABC) scale showed that healthy older people had more confidence undertaking activities without falling than people with PD (p < 0.04); the Ambulatory Self-Confidence Questionnaire (ASCQ) showed a similar trend but the group difference was not statistically significant (p = 0.07). It is worth noting that the ABC and ASCQ scores were also more variable for the PD than for the older group.

**Table 2.**
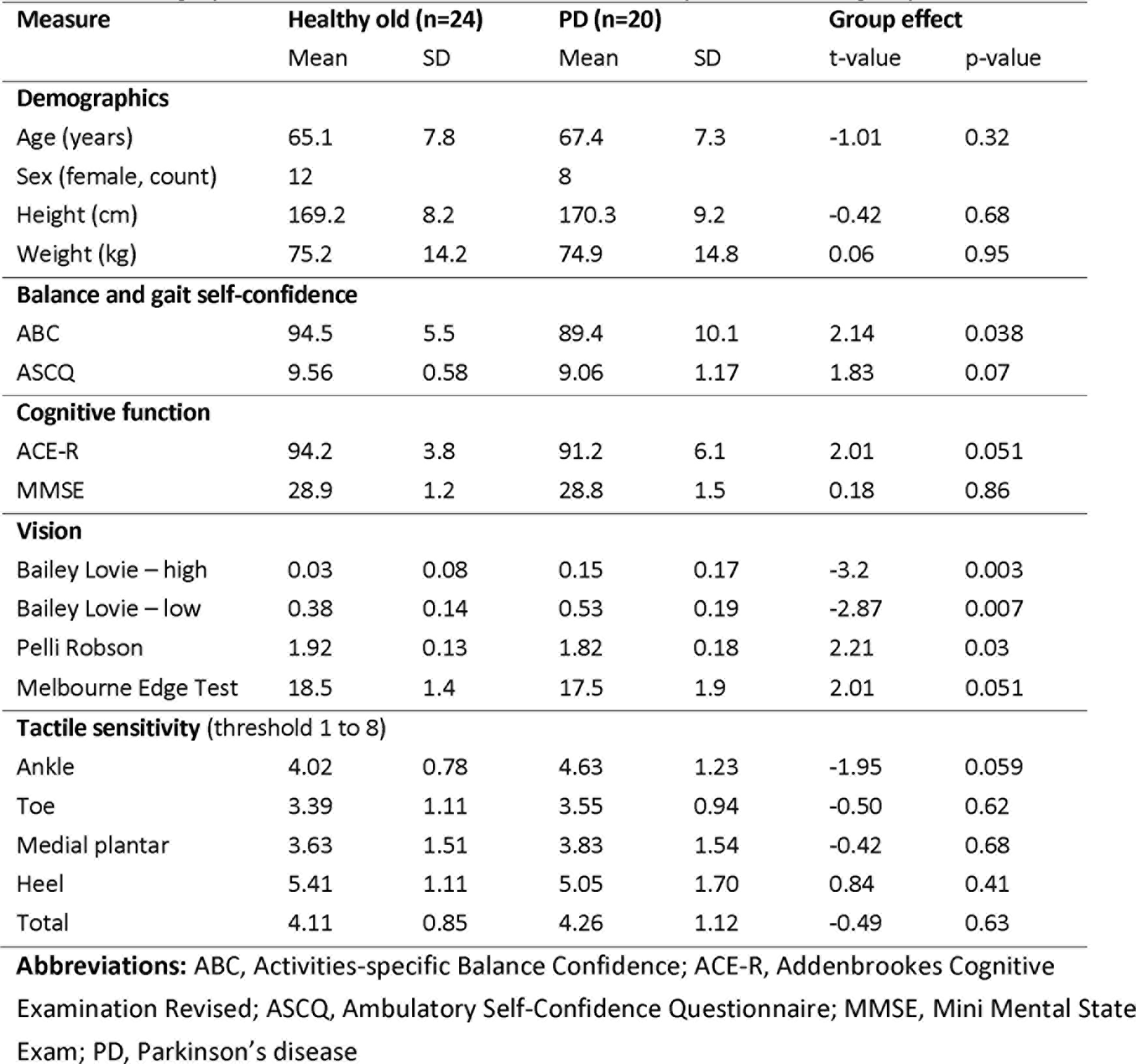
Demographics and clinical characteristics of healthy older and PD groups

### Temporal gait parameters

Most temporal gait parameters were similar for all groups (p ≥ 0.09) (Table 3). The duration of swing and single support was significantly different between groups (F_2, 63_ = 3.64, p = 0.03). Healthy older people had a significantly shorter swing and single support phase than healthy young people (p = 0.016), and so did people with PD (p = 0.027). There was no significant difference in swing or single support time between healthy older people and people with PD (p = 0.91). The main effect of condition (overground vs treadmill walking across all participants) for all gait parameters is presented in Table S1 in the Supplementary Material. Stride time, step time, stance phase and swing/single support phase were significantly shorter during treadmill compared to overground walking.

**Table 3.** Temporal Gait Parameters in healthy young, healthy older and PD groups

| Gait parameter | Healthy young |  | Healthy old |  | PD |  | Group effect |  |
| --- | --- | --- | --- | --- | --- | --- | --- | --- |
|  | mean | SEM | mean | SEM | mean | SEM | F | P |
| Walking speed (km/h) | 4.16 | 0.12 | 4.00 | 0.11 | 3.97 | 0.12 | 0.84 | 0.44 |
| Stride time (s) | 1.10 | 0.02 | 1.09 | 0.01 | 1.06 | 0.02 | 1.45 | 0.24 |
| Stride time variability (s) | 0.020 | 0.002 | 0.019 | 0.002 | 0.021 | 0.002 | 0.46 | 0.64 |
| Step time (s) | 0.548 | 0.008 | 0.541 | 0.008 | 0.522 | 0.009 | 2.48 | 0.09 |
| Step time variability (s) | 0.012 | 0.002 | 0.010 | 0.002 | 0.012 | 0.002 | 0.35 | 0.71 |
| Stance phase (s) | 0.689 | 0.013 | 0.696 | 0.012 | 0.674 | 0.014 | 0.74 | 0.48 |
| Swing phase/ Single support (s) | 0.404 | 0.007 | 0.381 | 0.006 | 0.382 | 0.007 | **3.79** | **0.03** |
| Double support (s) | 0.146 | 0.010 | 0.156 | 0.010 | 0.160 | 0.011 | 0.53 | 0.59 |

### Time-frequency power and coherence spectra

Event-related changes in time-frequency power and coherence within the gait cycle were assessed to investigate differences in neural control of gait between groups. This included estimates of five spectral measures using time-frequency decomposition: EEG and EMG power, inter-trial coherence of EMG and EEG, and corticomuscular coherence. These spectral estimates revealed a consistent pattern across groups: Power and coherence increased across a range of frequencies during double support and was reduced during the swing and single support phases. Figure 1 shows examples of time-frequency spectra of the right sensorimotor cortex and EMG from the left TA during overground walking in young, older and people with PD.

**Figure 1.**
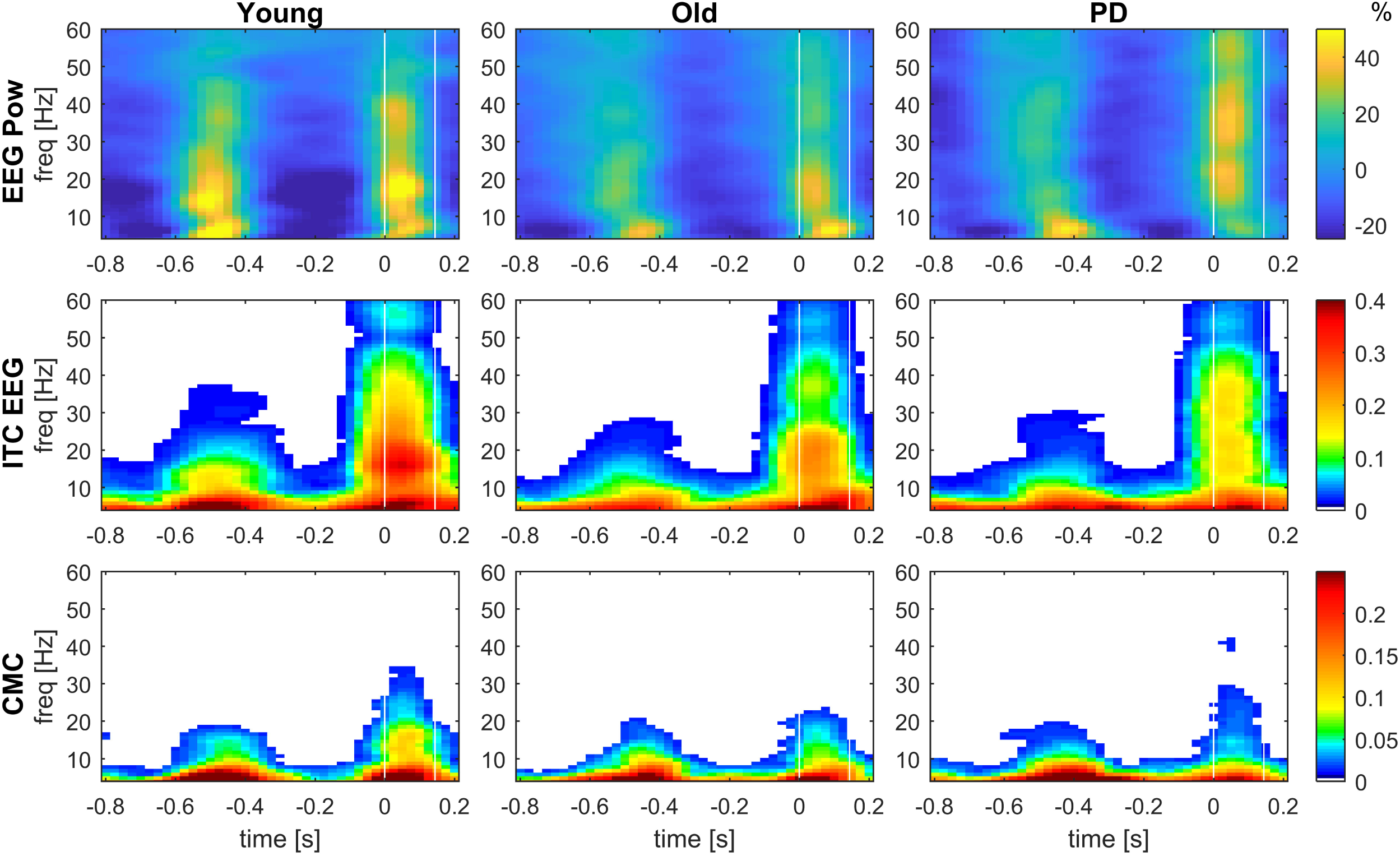
Grand-average time-frequency spectra of EEG power, inter-trial coherence and corticomuscular coherence. Event-related EEG power (EEG Pow, top row), inter-trial coherence of EEG (ITC EEG, middle row) and corticomuscular coherence (CMC, bottom row) acquired from bipolar EEG signals of the right sensorimotor cortex (C3-F3) and EMG from the left TA during overground walking in the young, old and PD groups. EEG power shows the percent change from the average. Coherence values (ITC and CMC) are thresholded: average coherence values below the 95% CI are set to zero (black). The x-axis shows the time in seconds relative to heel strike (t=0) of the left foot and the y-axis the frequencies in Hz. White horizontal lines indicate the double-support phase (from left heel strike to right toe off).

The averaged power and coherence across the double-support phase (0-125 ms after heel strike) are shown in Figure 2. Most of the power and coherence during the double-support phase was observed at low frequencies and decreased at higher frequencies. In addition, a peak around 16 Hz can be observed in EEG power and inter-trial coherence, which is most prominent in the young group. A peak in the low beta band (13-21 Hz) can also be observed in EMG power, inter-trial coherence and corticomuscular coherence, although these are less pronounced.

**Figure 2.**
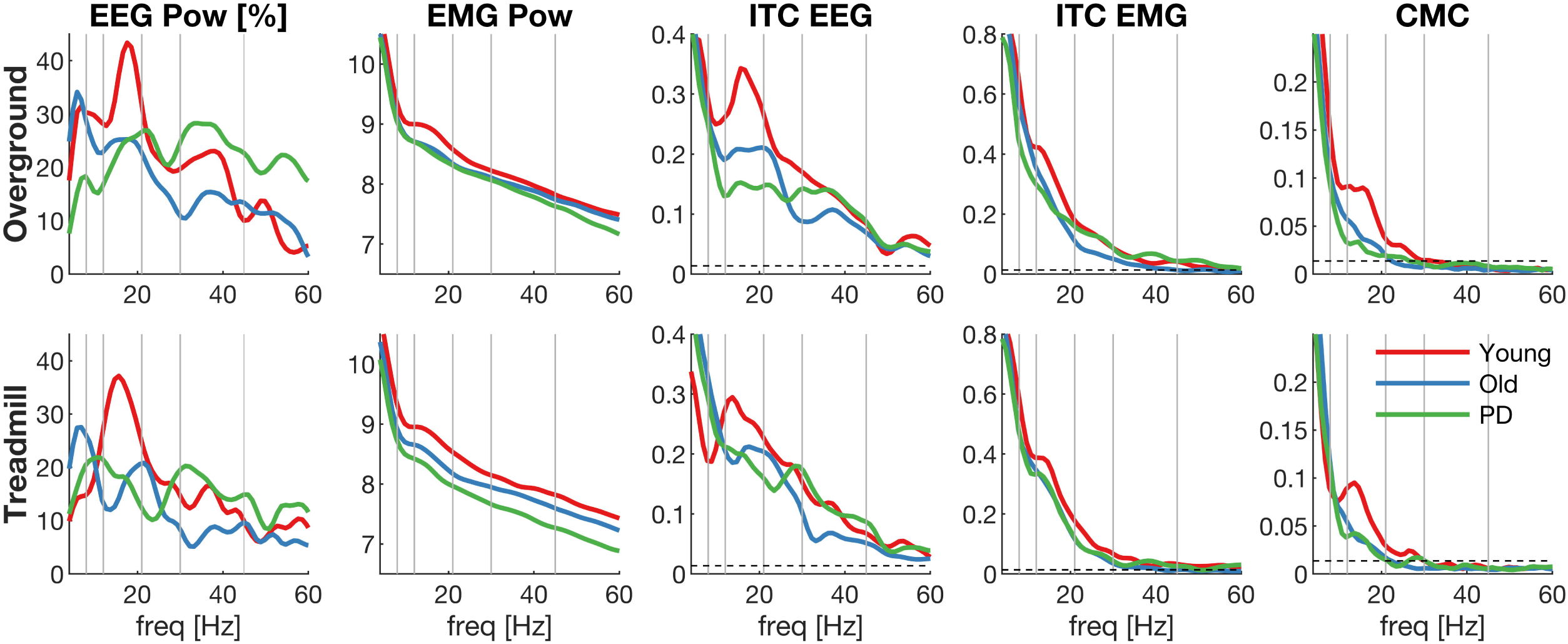
Grand-average frequency spectra during double support. Spectral measures averaged over double support for each group (young, old, PD): EEG power (first column), EMG power (second column), inter-trial coherence of EEG (ITC EEG, third column), inter-trial coherence of EMG (ITC EMG, forth column) and corticomuscular coherence (CMC, fifth column) during overground (top row) and treadmill (bottom row) walking. Data derived from bipolar EEG signals of the right sensorimotor cortex (C3-F3) and EMG from the left TA. X-axis shows frequencies in Hz; grey vertical lines highlight the different frequency bands and the dashed horizontal line the 95% CI of the coherence estimates.

The spectral measures were statistically compared between groups for five frequency bands (theta, alpha, low beta, high beta and gamma) using linear mixed models (LMMs) and the false discovery rate was controlled using the Benjamini-Hochberg procedure^96,97^. Table 4 reports the main effect of group for all spectral measures; supplementary figures S2 and S3 present these statistical results separately for overground and treadmill walking. Several significant main effects of group were observed during double support: EMG power at all frequency windows (p_adj_ < 0.02); EMG inter-trial coherence at theta frequencies (p_adj_ = 0.025); corticomuscular coherence at low beta frequencies (p_adj_ = 0.013). Similarly, several significant main effects of condition (overground, treadmill) were found (see supplementary Table S4) whereby magnitudes were reduced for treadmill walking: EEG power at theta, high beta and low gamma frequencies; EMG power across all frequency windows; EMG inter-trial coherence at high beta and gamma frequencies; corticomuscular coherence at high beta and gamma frequencies.

**Figure 3.**
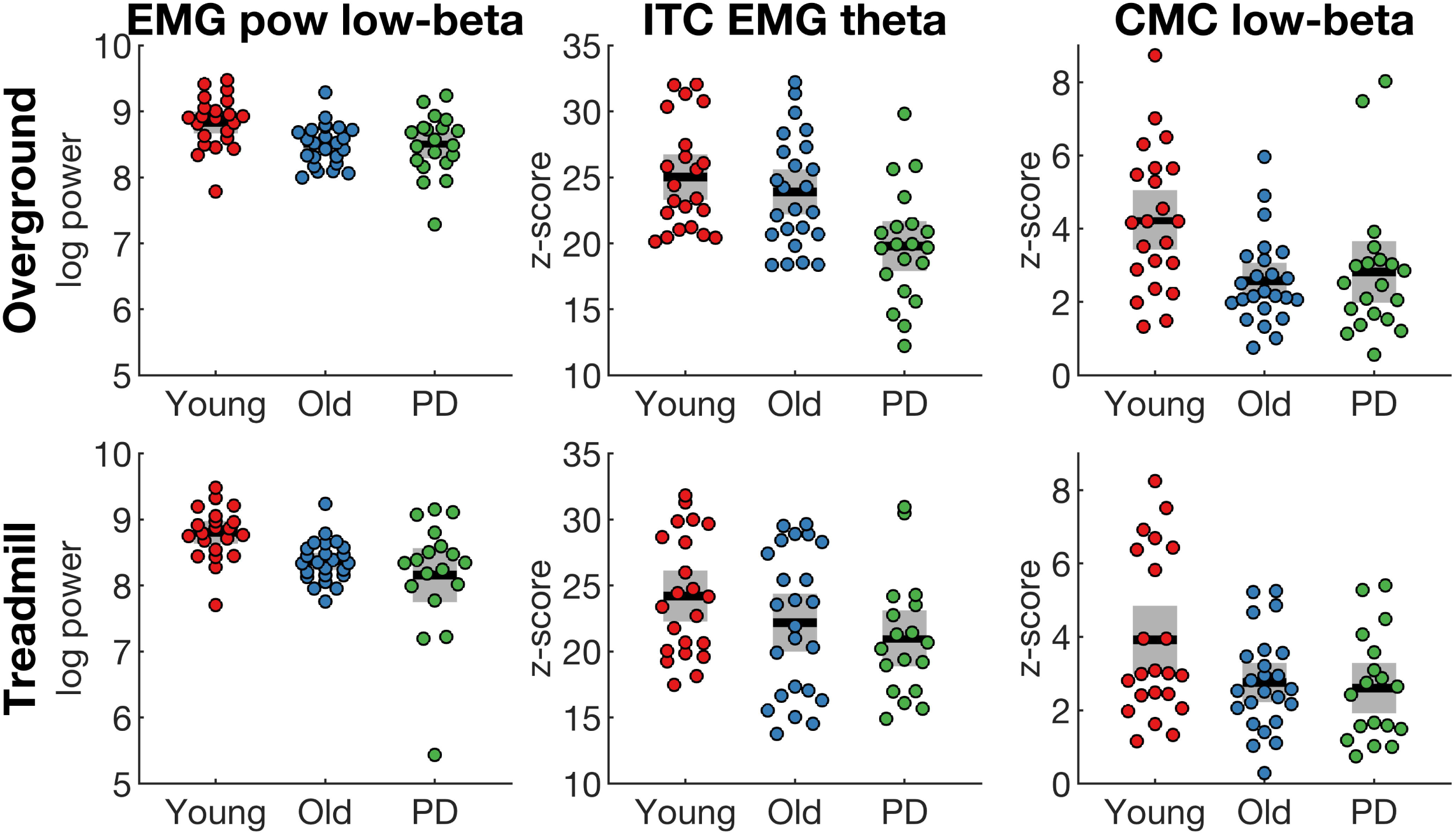
Spectral measures that reveal significant main effect of group. Spectral measures were averaged over double support in different frequency bands for each group (young, old, PD). EMG power at low beta frequencies (EMG pow, first column), EMG inter-trial coherence at theta frequencies (ITC EMG, second column), corticomuscular coherence at low beta frequencies (CMC, third column) during overground (top row) and treadmill (bottom row) walking. Data derived from bipolar EEG signals of the right sensorimotor cortex (C3-F3) and EMG from the left TA. Coloured dots show individual data of each participant, black horizontal lines show the group mean, grey boxes show the SEM.

**Table 4.** Main effect of group for spectral measures and mean estimates with 95% confidence intervals

| Spectral measure | Healthy young |  | Healthy old |  | PD |  | Group effect |  |  |
| --- | --- | --- | --- | --- | --- | --- | --- | --- | --- |
|  | mean | 95% CI | mean | 95% CI | mean | 95% CI | F | P | P<sub>adjusted</sub> |
| **EEG power (%)** |  |  |  |  |  |  |  |  |  |
| Theta | 21.7 | 14.4-29.0 | 23.6 | 16.6-30.56 | 16.1 | 8.3-23.9 | 1.05 | 0.36 | 0.50 |
| Alpha | 24.9 | 17.4-32.4 | 21.7 | 14.5-28.91 | 17.6 | 9.7-25.6 | 0.89 | 0.42 | 0.51 |
| Low beta | 33.2 | 25.2-41.2 | 20.0 | 12.4-27.68 | 19.7 | 11.2-28.2 | 3.71 | 0.030 | 0.075 |
| High beta | 20.6 | 13.2-28.1 | 15.8 | 8.7-22.94 | 18.9 | 11.0-26.9 | 0.45 | 0.64 | 0.67 |
| Gamma | 15.5 | 10.0-21.0 | 11.8 | 6.6-17.05 | 19.9 | 14.0-25.7 | 2.09 | 0.13 | 0.22 |
| **EMG power (log)** |  |  |  |  |  |  |  |  |  |
| Theta | 9.9 | 9.7-10.1 | 9.5 | 9.4-9.72 | 9.4 | 9.1-9.6 | **8.22** | **0.001** | **0.013** |
| Alpha | 9.1 | 8.9-9.2 | 8.7 | 8.6-8.9 | 8.6 | 8.4-8.8 | **5.86** | **0.005** | **0.021** |
| Low beta | 8.8 | 8.6-9.0 | 8.4 | 8.3-8.61 | 8.3 | 8.1-8.5 | **7.45** | **0.002** | **0.013** |
| High beta | 8.4 | 8.2-8.6 | 8.1 | 7.9-8.26 | 7.9 | 7.8-8.1 | **6.77** | **0.002** | **0.013** |
| Gamma | 8.0 | 7.9-8.2 | 7.8 | 7.7-7.98 | 7.6 | 7.4-7.8 | **6.16** | **0.004** | **0.020** |
| **ITC EEG (z-score)** |  |  |  |  |  |  |  |  |  |
| Theta | 12.5 | 11.1-13.8 | 13.0 | 11.7-14.29 | 12.3 | 10.8-13.8 | 0.28 | 0.76 | 0.76 |
| Alpha | 10.2 | 8.9-11.6 | 10.1 | 8.8-11.41 | 9.2 | 7.7-10.7 | 0.64 | 0.53 | 0.58 |
| Low beta | 11.1 | 9.7-12.6 | 9.1 | 7.7-10.49 | 8.1 | 6.6-9.7 | 4.20 | 0.019 | 0.060 |
| High beta | 8.7 | 7.2-10.1 | 7.6 | 6.2-8.97 | 7.2 | 5.6-8.7 | 1.10 | 0.34 | 0.50 |
| Gamma | 6.1 | 5.0-7.3 | 5.2 | 4.1-6.35 | 5.9 | 4.6-7.1 | 0.66 | 0.52 | 0.58 |
| **ITC EMG (z-score)** |  |  |  |  |  |  |  |  |  |
| Theta | 24.6 | 22.8-26.4 | 23.0 | 21.3-24.8 | 20.3 | 18.4-22.2 | **5.36** | **0.007** | **0.025** |
| Alpha | 15.9 | 14.5-17.4 | 14.5 | 13.0-15.88 | 13.0 | 11.5-14.6 | 3.60 | 0.033 | 0.075 |
| Low beta | 11.7 | 10.5-12.8 | 9.7 | 8.6-10.82 | 10.3 | 9.1-11.5 | 2.99 | 0.057 | 0.11 |
| High beta | 6.9 | 5.8-8.0 | 5.4 | 4.3-6.45 | 5.5 | 4.4-6.7 | 2.29 | 0.11 | 0.20 |
| Gamma | 3.6 | 2.9-4.4 | 3.0 | 2.3-3.68 | 3.1 | 2.3-3.9 | 0.89 | 0.42 | 0.51 |
| **CMC (z-score)** |  |  |  |  |  |  |  |  |  |
| Theta | 8.7 | 7.7-9.8 | 8.4 | 7.4-9.45 | 7.44 | 6.3-8.6 | 1.53 | 0.23 | 0.36 |
| Alpha | 5.1 | 4.4-5.9 | 4.4 | 3.7-5.16 | 3.75 | 2.9-4.6 | 3.02 | 0.056 | 0.11 |
| Low beta | 4.1 | 3.5-4.7 | 2.7 | 2.1-3.22 | 2.72 | 2.1-3.3 | **7.82** | **0.001** | **0.013** |
| High beta | 2.0 | 1.5-2.4 | 1.2 | 0.8-1.6 | 1.30 | 0.8-1.8 | 3.89 | 0.026 | 0.072 |
| Gamma | 0.9 | 0.7-1.1 | 0.7 | 0.5-0.9 | 0.74 | 0.5-1.0 | 0.87 | 0.43 | 0.51 |

CI, confidence interval; ITC, inter-trial coherence; CMC, corticomuscular coherence.

Post-hoc tests of significant group main effects revealed significant differences in EMG power and low-beta corticomuscular coherence between the young group and the old and PD groups but not between the old and PD groups (see Figure 3 and Table S5). In contrast, inter-trial coherence in the theta band was different between young and PD, as well as old and PD, but not between the young and old groups. Effect sizes (Cohen’s d_s_) of pairwise group comparisons are shown in Table S6.

### EMG envelopes

To further explore the group differences observed in spectral measures of EMG, the event-related EMG envelopes were analysed in the time domain. EMG envelopes revealed two bursts of activity during early swing and around heel strike (Figure 4). The second increase in EMG activity is marked by a double peak (at heel strike (t = 0) when the foot is fully dorsiflexed and at 60 ms after heel strike). Notably, the EMG amplitude was larger in older people and people with PD during early swing, and the second peak during the double support phase appeared to be reduced in the PD group. To statistically test this, we ran a LMM on the average EMG amplitude between −0.4 to −0.25 s (foot lift) and between 0.04 to 0.08 s (foot drop) relative to heel strike. EMG amplitude at foot lift was significantly different between groups (F_2, 63_ = 17.81, p < 0.0001), but not at foot drop (F_2, 63_ = 2.12, p = 0.13). Healthy older people had a significantly larger EMG amplitude during foot lift (67.3% [95%CI 62.6, 72.0]) than healthy young people (47.6% [95%CI 42.8, 52.5], p < 0.0001), and so did people with PD (61.9% [95%CI 56.7%, 67.0], p = 0.0001). There was no significant difference in EMG amplitude at foot lift between healthy older people and people with PD (p = 0.12). While no significant group effect was found during foot drop, we noticed considerable variability between participants in each group (Figure 5). For example, in the PD group EMG amplitude during foot drop varied almost between 5-90% relative to the maximum EMG amplitude within the gait cycle. The results of the main effect of condition are shown in Table S7, showing that the EMG amplitude during foot drop is lower during treadmill compared to overground walking.

**Figure 4.**
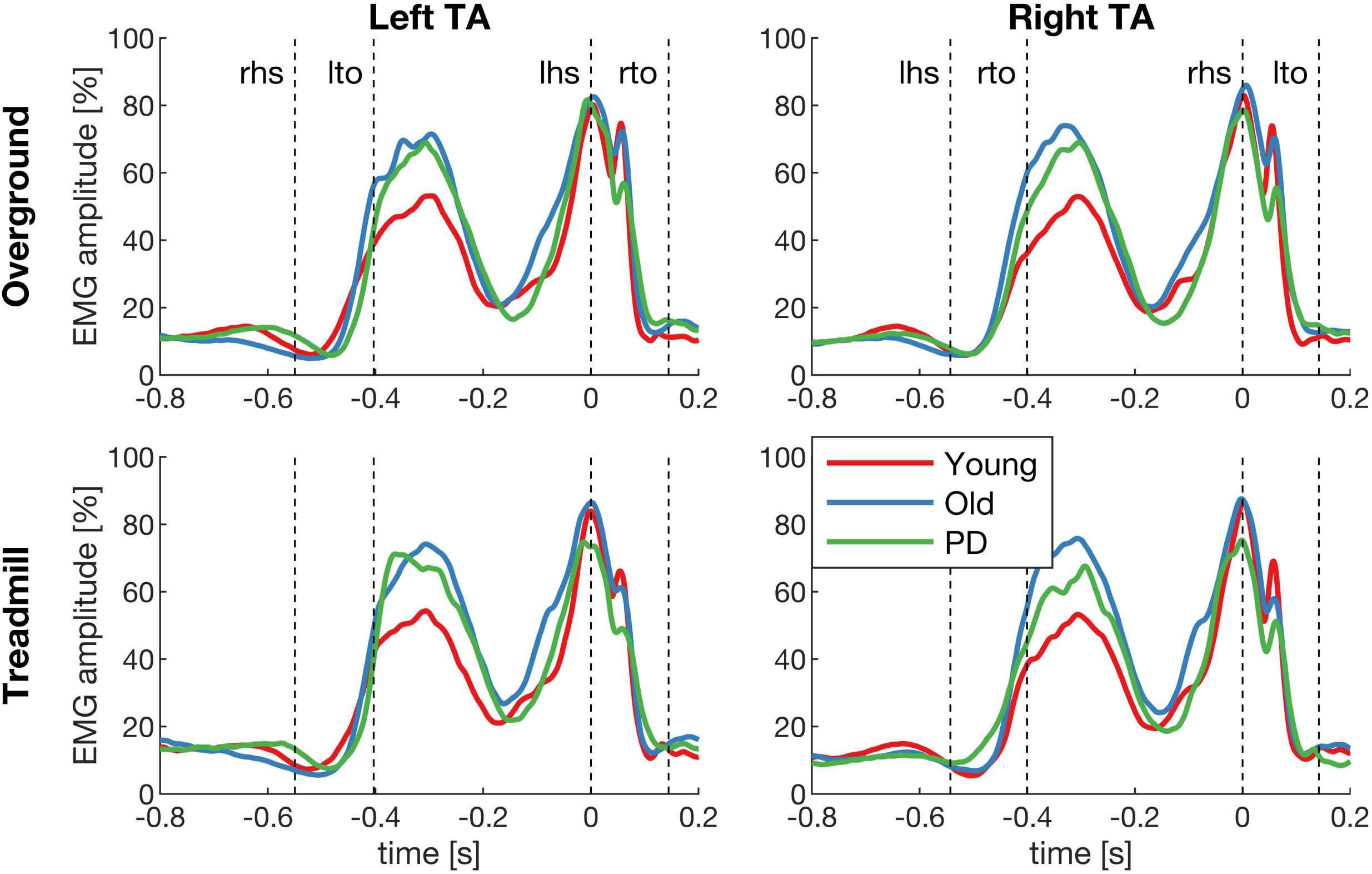
Grand-average EMG envelopes. EMG envelopes of the left and right tibialis anterior (TA) during overground and treadmill walking in the young, old and PD groups. The x-axis shows time relative to heel strike (t=0) and the y-axis the amplitude relative to the maximum within the gait cycle. Dashed lines show the kinematic events (lhs, left heel strike; lto, left toe-off; rhs, right heel strike; rto, right toe-off).

**Figure 5.**
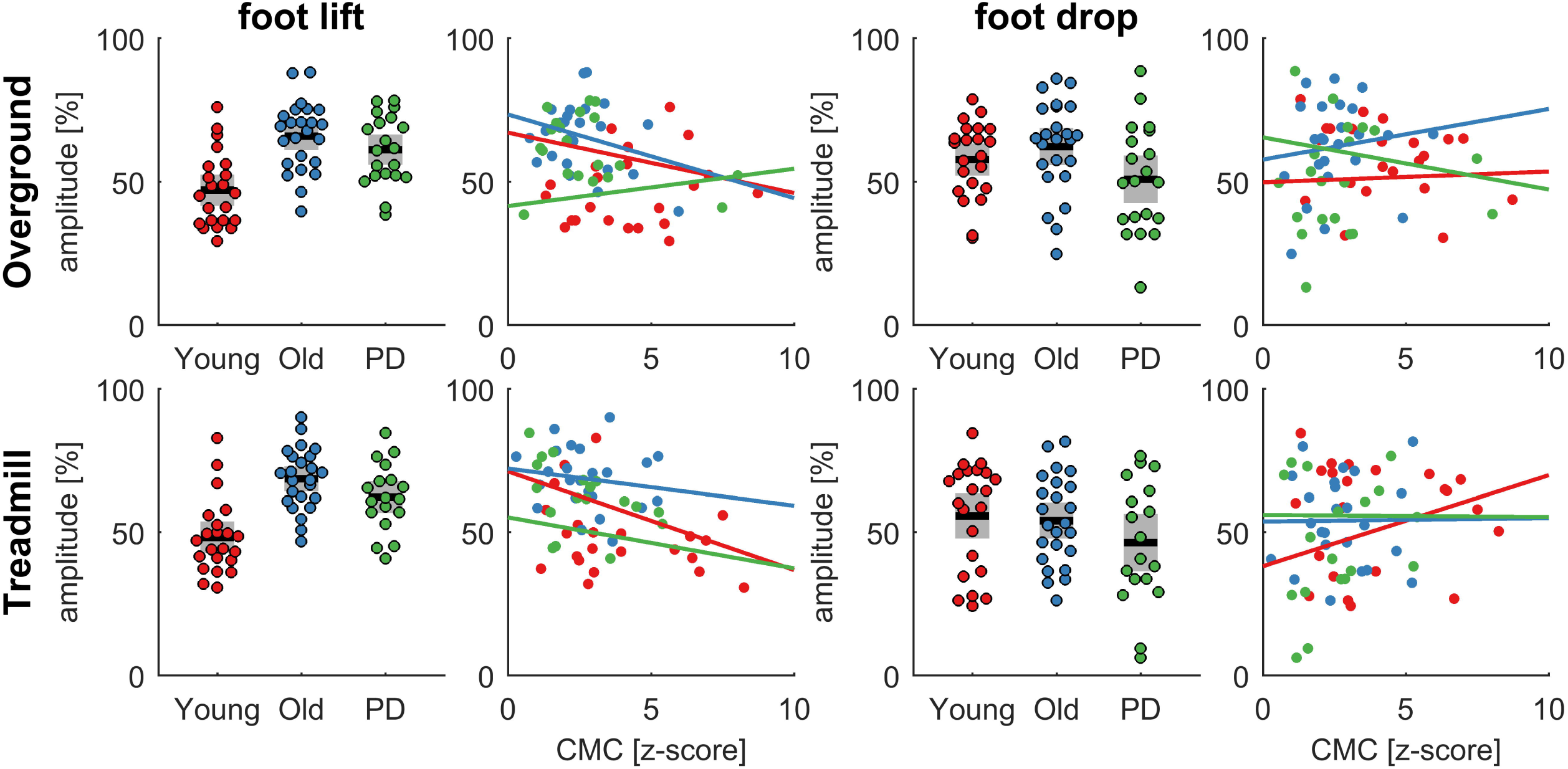
Relationship between EMG amplitude and corticomuscular coherence. EMG amplitude was averaged in two time intervals (foot lift: −0.4 to −0.25 s; foot drop: 0.04 to 0.08 s for each group (young, old, PD) and condition (overground and treadmill). The bar graphs show the summary statistics in each group and the scatter plots the relationship with corticomuscular coherence (CMC) in the lower beta band. Coloured dots show individual data of each participant, black horizontal lines show the group mean, grey boxes show the SEM.

The EMG amplitude during both time intervals was correlated with corticomuscular coherence in the lower beta band during the double support phase within each of the groups (young, old and PD) and conditions (treadmill and overground). None of the correlations were significant when testing each group separately (p > 0.05). EMG amplitude during foot lift appears to be negatively correlated with corticomuscular coherence (Figure 5, left panels), but this is mainly due to the group effects in which the young group shows lower EMG amplitude during foot lift and higher corticomuscular coherence during double support.

## Discussion

We performed time-frequency analysis of electrophysiological data recorded during overground and treadmill walking in healthy young, healthy older and people with PD to investigate changes in cortical and corticospinal processes during walking. PD participants were in early stages of the disease with mild to moderate symptom severity, without freezing of gait or frequent falls and were optimally medicated for all assessments. All three groups showed significant corticomuscular coherence at theta, alpha, beta and gamma frequencies (4-45 Hz) between the sensorimotor cortex and the contralateral TA muscle during the double support phase that was largely absent during the swing phase. Similarly, power and inter-trial coherence also increased during double support. Older participants and people with PD showed significantly reduced corticomuscular coherence at low beta frequencies (13-21 Hz) compared to young individuals, but there was no difference between healthy older and people with PD. EMG power was significantly decreased in all frequency bands in healthy older and PD participants compared to the healthy young group. Interestingly, walking speed, stride time and its variability, and the duration of the double support phase was similar across all groups. However, people with PD and healthy older people spent significantly shorter time in single support and swing phases of gait than healthy young people. The amplitude of the TA EMG envelope during early swing was significantly larger in older people and people with PD compared with healthy young participants. Hence, the current findings reveal age-related changes in the corticomuscular control of walking. They further indicate that corticomuscular control in people with early-stage PD and low to moderate disease severity while under optimal medication is similar to that of older people. Task-dependent differences in neural control of locomotion are also suggested by temporal gait and frequency-dependent differences between overground and treadmill walking.

### Changes in corticomuscular control with aging and Parkinson’s disease

We observed several differences in electrophysiological and gait measures between young and old participants, as well as young and PD participants, but not between the old and PD groups. A primary outcome was reduced corticomuscular coherence at low beta frequencies during double support for older and PD participants compared to young participants. This indicates that reduced functional connectivity in corticospinal pathways involved in gait control is primarily affected by age. Several previous studies also found reduced corticomuscular coherence in older compared to younger adults. Spedden et al.^98^ reported attenuated corticomuscular coherence during treadmill walking in older compared to young participants. Yoshida and colleagues^82,83^ found decreased corticomuscular coherence during rhythmic ankle movements in older people compared to young adults but not between older and PD participants, and Ozdemir et al.^99^ found reduced corticomuscular coherence for the TA muscle during perturbed standing in older compared to young people. Studies on upper limb muscles during sustained contractions also found reduced corticomuscular coherence in older people^85,86^. However, other studies on forearm and finger contractions found increased corticomuscular coherence in older adults^84,100^, which may reflect task-specific requirements (visual feedback, dual-tasking). In people with PD, corticomuscular coherence during tonic forearm contractions was also not found to be different compared to healthy age-matched controls^87,88^, and may^87^ or may not^88,101^ be modulated by levodopa medication. As PD participants in this study had a low to moderate disease severity, similar levels of cognitive function, tactile sensitivity and gait as their healthy age-matched counterparts, and they completed the experiments on medication, this may explain why we did not observe a difference in corticomuscular coherence between PD and age-matched controls.

We did not find a significant group effect for EEG power and EEG inter-trial coherence, at least not after controlling the false discovery rate (see Table 4). However, the direction of the observed differences between groups was still similar to the changes in corticomuscular coherence, especially at low beta frequencies (see Figure 2). This may imply that a group difference was present but that the effect size was too small to detect it with the current sample size. Possibly, measures based on EEG alone are less sensitive (higher inter-subject variability) than measures based on EEG and EMG (i.e. corticomuscular coherence). Previous studies have documented changes in cortical oscillatory activity with age and PD mainly during fine motor tasks of the upper limbs and at rest^63,102–106^. During walking, EEG gamma power over frontal cortices was enhanced in older people during dual-task walking compared to young controls^59^. During upright standing, cortical activity was altered in older people^99,107,108^. In people with PD, increased theta and low-gamma cortical power was associated with freezing of gait during turning compared to normal fluent turning^60,61^. However, whether cortical oscillations during walking are truly affected by age or PD remains to be shown in larger future studies.

In addition to the changes in the corticomuscular coherence results, we also found reduced EMG power during double support in older and PD participants compared to young participants. Again, no significant differences were observed between the older and PD groups. Previous studies reported a number of age-related changes in EMG activity during walking indicating that recruitment of ankle muscles changes with age. For example, EMG magnitude of the TA at mid swing was shown to decrease with age^109^, and the onset of TA activity within the gait cycle was earlier and offset was later in older compared to young adults^110,111^. In PD, EMG activity of distal leg muscles during gait was found to be reduced and poorly modulated^112–115^. As motor units undergo several morphological and physiological adaptations with ageing, this also affects their discharge behaviour^116^. For instance, motor unit firing rates decrease with age^117–119^ and show greater variability^120^. We also found a significant difference in inter-trial coherence of the EMG signal at theta frequencies between groups, but unlike the other outcome measures it was reduced in people with PD compared to young and old controls (Table S5). This suggests reduced phase-locking of the EMG envelopes with respect to heel strike, at least at theta frequencies, and may indicate more variable muscle activation patterns across gait cycles in people with PD.

Although EMG power was reduced during the double-support phase, we did not find a concomitant reduction in EMG amplitude in the time domain (see Figure 4). This may be partly due to the way EMG amplitude was normalised^121^. However, EMG envelope amplitude was significantly increased during early swing in older and PD compared to young people. Although the temporal profiles of the EMG envelopes in the present study are consistent with previous reports^113,122–126^, the direction of reported changes in EMG amplitude with age are only partially consistent. For example, some studies found increased EMG amplitude in older people during treadmill^125^ and overground walking^126^ at different speeds (in agreement with our study), while other studies reported a reduction in EMG amplitude with age during overground walking^124^. Likewise, age-related changes in EMG amplitude have been found in different parts of the gait cycle, e.g. at mid swing^124^, mid stance^126^, and terminal stance^124^. This may be due to differences in experimental designs (e.g. number and duration of walking trials and speeds), and due to data processing techniques (e.g. how EMG signals were normalised). Notably, we found the TA EMG amplitude during foot drop reduced during treadmill compared to overground walking in all subjects, which is in agreement with previous reports of lower TA activation during the load phase of the gait cycle in people with and without PD^33^.

Older and PD participants spent shorter time in swing and single support than young individuals indicating age-related changes in motor performance. This is consistent with previous reports of kinematic and kinetic changes of gait in older people^9,11^. Previous studies have also documented differences in gait between people with PD and healthy age-matched controls^1, 12–16, 112–114^, which we did not observe in the current study. However, previous studies mainly detected PD-related gait differences by kinetic, kinematic, spatial and muscle activation parameters but less consistently in temporal gait parameters, which were assessed in the current study. Although we did not find any differences in gait performance between people with PD and their healthy age-matched controls, we found differences in gait performance as well as in a number of spectral outcome measures and EMG envelopes between overground and treadmill walking. Treadmill walking reduced the duration of stride and step time, stance and swing phase, and the amplitude of the TA EMG at foot drop. The backward movement of the treadmill belt may shorten the time between heel strike and forefoot drop, which is largely controlled by the dorsiflexors and ensures body weight support during the early load phase^33,127,128^. Multiple spectral measures also showed reduced magnitudes during treadmill walking, in particular EEG and EMG power, EMG inter-trial coherence and corticomuscular coherence at high beta and low gamma frequencies. These differences in power and coherence may reflect changes in gait stability^129^, step adaptations^130^ or sensorimotor processing^91^ related to controlling foot drop^80^. Possibly, these mechanisms help to explain improvements in gait in people with PD after treadmill training^30,32–34^.

In summary, there were several age-related changes in physiological and temporal gait variables during overground and treadmill walking, but little or no differences between the older and PD groups. It is possible that the lack of group differences between older and PD participants in our study is due to characteristics of our particular PD sample (low to moderate disease severity, similar levels of cognitive function and tactile sensitivity as healthy controls, optimally medicated) or due to low statistical power of this study. The effect size estimates for the old-PD comparisons were mostly negative and on average |0.24|. It is hence possible that small differences exist between people with PD and age-matched controls which cannot be detected with the current sample size. The effect sizes for the comparison of the young and old groups were also mostly negative and many were larger than |-0.5|; the young-PD comparison effect sizes showed a similar trend with even larger values and a number of them reaching approximately |-1|. While these are rather large effect sizes, this should be interpreted with caution as effect sizes are often overestimated in small samples^131^. Indeed, the confidence intervals we report indicate that the true population effect sizes may be much smaller.

### Physiology of low beta oscillations

During the double support phase, the frequency spectra showed a gradual reduction in power and coherence at higher frequencies with a superimposed peak in the lower beta range (see Figure 2). The peak in the beta band was more pronounced in the EEG measures (EEG power and inter-trial coherence) than in the EMG measures. In this frequency band we also found a significant group effect for corticomuscular coherence and EMG power. The other outcome measures show a similar pattern in the low beta band (highest in young, decreased in older and PD participants; Figure 2), although these did not reach statistical significance in the group comparison. Notably, the effect size estimates in the low beta band were consistently negative across all outcome measures and largest for corticomuscular coherence and EMG power.

Oscillatory activity at beta frequencies has been identified as an important feature in motor control. Many studies have shown that beta band activity is enhanced during tonic contractions and diminished during dynamic voluntary movement^132–138^, and can be observed at different levels of the central nervous system including motor cortex, basal ganglia and motor units in the periphery^62,132^. Recently it was shown that movement-related modulations of beta oscillatory activity occur in brief bursts rather than as an ongoing rhythm^139,140^. Beta band activity is thought to reflect a mechanism involved in maintaining the current sensorimotor state^67,132,141^. During walking, beta oscillations have been found to be task-dependent and a number of interpretations have been put forward. For instance, cortical beta oscillations may be related to controlling gait stability^129,142^, control of step lengthening and shortening adaptations^130,143^, visuomotor integration^144–146^, speed control^147,148^, forward propulsion driven by ankle plantar flexors^149^, erroneous anticipatory postural adjustments during gait initiation^150^, and responding to sensorimotor conflict during perturbations while walking^151^. An interesting hypothesis is that activity in the beta band may underpin the temporal coordination of sensorimotor processes^152–154^, and hence be involved in the temporal coordination of the heel strike during walking.

In PD, it is well established that beta activity in the basal ganglia is enhanced at rest in the dopamine-depleted state and can be restored to normal levels with levodopa medication^63,64,155,156^. This exaggerated pathophysiological beta activity in the basal ganglia has been found associated with bradykinesia and impaired motor function^62–64,66,157^. Interestingly, walking and cycling has been found to modulate/normalise beta activity in the subthalamic nucleus (STN) in people with PD^73–75^. Walking and cycling was found to decrease STN beta power in people with PD^73^. Stepping movements were found to modulate STN beta power relative to the movement cycle, and these modulations were associated with stepping performance^74^. Patterns of STN beta oscillations (i.e. duration of beta bursts) were also found to be linked to freezing of gait^158^. A number of studies that investigated cortical oscillatory activity during upper limb movements reported beta frequencies to be affected by PD^88,101, 159–161^. For instance, cortical beta power during static forearm contractions was increased in people with PD compared with healthy controls^88^, although others reported it to be decreased^159,161^.

We only observed a reduction in low beta corticomuscular coherence during the double support phase. The duration of the double support phase is about 125 ms and fits about 2 cycles of beta oscillations. Hence, these results likely reflect reduced bursts of beta oscillations rather than changes in ongoing beta oscillations^162,163^. These beta bursts may represent brief activity reflecting the retrospective evaluation of task performance and updating subsequent task performance^139^. In the present context, error-related activity may reflect the temporal parameters of step cycles. We found a parallel increase in power and inter-trial coherence, which suggests that transient beta activity likely represents an evoked response rather than true synchronisation of two interacting oscillators^80^. As we observe coherent beta activity at the cortical and spinal level, it is possible that these beta bursts are transmitted via the corticospinal tract or that subcortical or spinal networks simultaneously evoke time-locked responses at the cortical and spinal level. As such, the decreased low beta corticomuscular coherence in older and PD groups found in this study may reflect an attenuated evoked response at cortical and/or spinal levels. At the spinal level, differences in evoked TA responses may reflect reduced activity of dorsal horn spinal neurons^164,165^, which receive input from Ia afferents that are known to degenerate with age. During upright standing, for example, age-related degeneration of Ia afferents has been shown to affect leg muscle activity and to contribute to impaired postural control performance^166^.

### Limitations

The current study is exploratory in nature involving five different outcome measures in five frequency bands (resulting in 25 statistical comparisons). With a limited samples size (about 20 per group) the statistical power is fairly low and results should be interpreted cautiously. We did control the false discovery rate using the Benjamini-Hochberg procedure (see adjusted p-values in Table 4). Still, future studies (that are ideally pre-registered)^167^ will need to confirm these findings by specifically testing in the lower beta band, which will increase statistical power.

Another limitation is that PD participants completed all experiments in an optimally medicated state and we therefore cannot address potential medication effects on gait and cortical/corticospinal oscillatory activity in people with PD. Past studies reported inconsistent and contradictory findings with respect to medication effects on cortical and corticospinal oscillations at rest and during movement in people with PD. For instance, Salenius et al.^87^ reported a levodopa effect on beta corticomuscular coherence during isometric forearm contractions, while Pollok et al.^88^ and Hirschmann et al.^101^ did not. Notably, Yoshida and colleagues^82,83^ found decreased corticomuscular coherence in older people compared to young adults during rhythmic ankle movements but not between older and PD participants (similar to our results) when tested their PD participants in an ‘off-medication state’ (after overnight withdrawal). This creates uncertainty with regards to potential medication effects on cortical and corticospinal oscillatory activity during walking in people with PD, and it is possible that we did not find differences between the PD and age-matched control group because the PD group was assessed in an on-medication state. Another potential explanation may be the characteristics of our particular PD sample (low disease severity, cognitive function, tactile sensitivity, ‘agility’), indicating that our PD sample may not have been very different from the healthy older cohort. Moreover, the experimental paradigm itself was not very challenging (simple steady-state walking) and may therefore not be able to distinguish minor deficiencies in corticomuscular control in this relatively agile sample. More challenging paradigms including dual-task walking or perturbations that trigger deliberate stepping adaptations (for example, similar to Malcolm et al.^57^) may be better suited to detect changes in cortical and corticospinal oscillatory activity during the early stages of PD.

Finally, mobile EEG is susceptible to movement artefacts and caution should hence be exerted when interpreting phasic changes in EEG activity^168–175^. In this study, we applied multiple strategies to minimize the effects of motion artefacts according to standards currently established in this research field. We excluded gait cycles with excessive artefacts, we performed ICA, and we calculated a bipolar derivative EEG signal^80^. The limited number of EEG channels recorded in this study may have compromised the source separation of artefact sources, however even acquisition of a larger number of EEG channels cannot guarantee that EEG signals will be artefact free^169^. We also pre-processed the EMG recordings by high-pass filtering, rectifying, and demodulating the signals in order to remove low-frequency artefacts and periodic amplitude modulations that could distort the coherence estimates^176,177^. Previous studies found that low delta and high gamma oscillatory activity may be more affected by movement artefacts than other frequency bands^168^. Notably, the time-frequency profiles observed in this study are similar to those in multiple previous ambulatory EEG studies^80,81,90,129,147, 178–181^. This suggests that the observed time-frequency profiles in the current and previous studies reflect neural activity, or alternatively, are all affected in a similar way by movement artefacts. Importantly, in the current study we found a significant group effect for corticomuscular coherence specifically at lower beta frequencies, but not over a broadband frequency range. It seems unlikely that these frequency-specific changes are caused by non-physiologic artefacts. Mobile EEG recordings remain challenging and methods to reduce movement artefact contamination (e.g. artefact subspace reconstruction^182^ or dual-electrode arrays^183^) are constantly being developed^170,184^.

### Conclusion

This study shows coherent corticomuscular activity during the double support phase of the gait cycle during steady-state overground and treadmill walking in healthy young and older adults, as well as people with PD. We found that low beta corticomuscular coherence and EMG power was decreased in older and PD participants compared to young people, and that it was not different between older and PD participants, suggesting that these alterations in corticomuscular control of gait were mainly age-related. Furthermore, task-dependent differences in neural control of locomotion were also suggested by frequency-dependent differences between overground and treadmill walking. Transient and concomitant increases in power and inter-trial coherence suggest evoked bursts of beta activity at spinal and cortical populations rather than a modulation of ongoing corticospinal beta synchronization. The reduction of transient beta activity may reflect changes in the temporal coordination of motor events within the gait cycle such as the timing of foot drop. Observed group differences in several electrophysiological and temporal gait measures indicate inter-related control mechanisms at the peripheral, spinal and cortical levels. However, as we did not observe any correlations between corticomuscular coherence and EMG amplitude (see Figure 5), this suggests that ageing- and PD-related changes are multi-factorial, i.e. affect different systems, including the cortex, spine and periphery.

## Materials and Methods

Three groups participated in the study with 69 participants in total: 1) 24 healthy young adults, 2) 24 healthy older adults, 3) 21 individuals with PD. All experimental protocols were approved by the Human Research Ethics Committee of Queensland University of Technology (#1300000579) in accordance with the Declaration of Helsinki, and all participants gave written informed consent prior to participation. The results of the young adults have been previously described in Roeder et al.^80^.

### Experimental protocol

Participants performed overground walking for 12-14 minutes and treadmill walking for approximately seven minutes. Participants walked barefoot at their preferred speed (3.3 – 4.8 km h^−1^), which was matched across conditions on an individual basis. During both conditions, participants walked with their hands free (natural arm swing). In the overground condition, participants walked back and forth along a straight path (∼14 m) on a firm surface in the gait laboratory and turned at each end of the room. The turning sections, including acceleration at the beginning of the straight-line path and deceleration at the end (∼2.4 m), were excluded from further analyses and only straight-line walking (8.9 m) was used for further analysis. The order of treadmill and overground walking was randomised across participants, and participants rested for 5 minutes between both walking conditions. Before the actual gait experiments commenced, participants completed a test trial (overground) of approximately three minutes in order to determine their preferred walking speed. Their mean walking velocity during the test trial was used to set the treadmill belt speed and to monitor and maintain the participants’ walking speed during the overground condition. If the participant did not comply with the pre-determined mean walking speed during the overground condition, they were instructed to walk faster/slower.

Healthy older people and people with PD also completed the following tests on a day other than the walking experiments within a 6-week period: Addenbrooke’s Cognitive Examination (ACE-R) and Mini-Mental-State-Exam (MMSE)^94,185^, visual function tests (Bailey Lovie, Melbourne Edge Test, Pelli Robson)^186–188^, and a peripheral sensation test assessing tactile sensitivity at different sites of their feet (lateral malleolus, plantar surfaces of the great toe, midfoot/arch and heel) using Semmes-Weinstein pressure aesthesiometer with nylon monofilaments^95^. Moreover, participants completed a number of health-related questionnaires including the ABC scale^189^, ASCQ^190^, and Edinburgh Handedness Inventory ^191^. Additionally, people with PD were assessed for disease severity using the Unified Parkinson’s Disease Rating Scale (MDS-UPDRS) and Hoehn & Yahr scoring^192,193^, and filled in the PD Gait and Falls questionnaire^194^. People with PD presented in their optimally medicated state for all experiments.

### Data acquisition gait experiments

EEG, EMG and kinematic events (heel strike and toe off) were recorded while participants performed both gait conditions (overground and treadmill). Bipolar surface EMG was recorded from the left and right tibialis anterior muscle (TA) and filtered at 1-1000 Hz. Simultaneously, EEG signals were recorded using water-based AgCl electrodes placed at 10 cortical sites according to the international 10-20 standard (P3, P4, C3, Cz, C4, F7, F3, Fz, F4, F8), and filtered at 1-500 Hz. Footswitches were attached onto the participants’ sole at the heel and the big toe of both feet. All signals (EEG, EMG and footswitches) were recorded synchronously with a wireless 32-channel amplifier system (TMSi Mobita, The Netherlands) and sampled at 2 kHz. The recording system and wireless transmitter were placed in a belt bag and tied around the participants’ waist.

### Data analysis

All processing of EEG, EMG and footswitch data were performed in MATLAB (2017a) using custom written routines.

Prior to spectral analysis, electrophysiological data were normalized to unit variance and pre-processed to remove artefacts. EEG channels were first visually inspected and segments with excessive noise (large-amplitude movement artefacts, EMG activity) were removed. EEG signals were then band-pass filtered (2^nd^ order Butterworth, 0.5-70 Hz) and re-referenced to a common average reference (EEGLAB version 13.6.5)^195^, as recommended in Snyder et al.^170^. Subsequently, independent component analysis (ICA) was performed using the infomax ICA algorithm implemented in EEGLAB. Independent components containing eye blink, muscle, or movement artefacts were removed from the data. On average, 3.8 components (SD 0.9) were removed from participants in the healthy young group, 3.9 (SD 0.9) from healthy older participants, and 4.2 (SD 1.0) from participants with PD. The remaining components were retained and projected back onto the channels. Finally, we calculated bipolar EEG montages based on the monopolar recordings to assess cortical activity from bilateral sensorimotor cortices: C3-F3 for the left sensorimotor cortex, and C4-F4 for the right sensorimotor cortex^196^. A differential recording between electrode pairs suppresses far-field contamination and is hence most sensitive to local activity generated between the electrode pairs. Thus, it can be used to attenuate artefacts similar to a common reference approach (e.g. Snyder et al. 2015^170^, Petersen et al. 2012^79^). An example of the processing steps of the EEG signals of one healthy older participant during overground walking is shown in the supplementary figure S8.

EMG data were high-pass filtered (4^th^ order Butterworth, 20 Hz cut-off) and full-wave rectified using the Hilbert transform^197^. For coherence analysis EMG signals were also demodulated (the Hilbert amplitude was normalized while leaving the Hilbert phase unchanged) to avoid spurious coherence estimates resulting from periodic changes in EMG amplitude^176^.

### Time-frequency analysis

We then used time-frequency analysis to assess changes in spectral measures within the gait cycle. To this end, EEG and EMG signals were segmented into 220 segments of 1-s length (−800 to +200 ms with respect to heel strike). Heel strike served as reference point (t = 0) in the gait cycle to which all segments were aligned. Event-related power and coherence spectra were estimated across the 220 segments using Short-time Fourier transform (a 375-ms Hanning window with increments of 25 ms). We computed five spectral measures: EEG and EMG power, inter-trial coherence of EEG and EMG signals, and corticomuscular coherence^198^. Event-related EEG power was expressed as percentage change from the average by subtracting and dividing by the mean power across the time interval separately for each frequency^199^. Here we analysed changes in relative power and hence focused on phasic changes in power as tonic differences were subtracted away. Event-related EMG power was log transformed. EMG power was analysed to assess common input to motor neuron pool of the TA at these frequencies^200^. Event-related power and coherence spectra were computed for each participant during overground and treadmill walking, and subsequently averaged across participants to render the grand-average for each estimate and condition. Additionally, event-related EMG envelopes were computed using filtered and rectified EMG data. The EMG envelopes were smoothed (2th order low-pass filter at 45 Hz) and expressed as a percentage of the peak amplitude within the gait cycle^201^.

### Statistical analysis

The significance level (alpha) was set at 0.05 for all statistical analyses, which were performed in SPSS (version 25). Linear mixed models (LMM) were used to compare gait parameters across groups. Fixed effect factors included in the model were group (young, old and PD) and condition (overground, treadmill). The random effect was participant. When a significant main effect was found for a comparison, Fisher’s LSD was used as post-hoc test.

Clinical measures available for the old and PD groups (ACE-R, MMSE, visual function measures, tactile sensitivity, ABC, ASCQ) were compared between groups via independent samples t-tests. Disease characteristics of PD such as MDS-UPDRS scores, Hoehn & Yahr (H&Y), Gait and Falls questionnaire (GFQ) and levodopa equivalent daily dosage (LEDD) were summarised as group means.

Coherence (corticomuscular coherence and inter-trial coherence) was transformed to z-scores for statistical comparison between groups and conditions. A parametric approach was used to convert magnitude-squared coherence to p-values^202^, and we then used the standard normal distribution to convert p-values into z-scores^80^, which have an expected value of 0 and a standard deviation of 1 under the null hypothesis (EEG and EMG signals are independent).

We used LMMs again to compare spectral measures across groups and conditions. Spectral estimates (EEG power, EMG power, EEG inter-trial coherence, EMG inter-trial coherence, corticomuscular coherence) were assessed in five time-frequency windows of interest: Double support (0-125 ms relative to heel strike) at theta (4-7 Hz), alpha (8-12 Hz), lower beta (13-20 Hz), higher beta (21-30 Hz), and gamma frequencies (31-45 Hz). Hence, 25 separate LMMs were completed (one for each outcome measure and frequency band), and the false discovery rate was controlled using the Benjamini-Hochberg procedure^96^. We used a Matlab script to control the false discovery rate^97^. The fixed effect factors included in the LMMs were group and condition, and the random effect was participant (i.e. using the participant identification number), which accounts for the repeated spectral measures. Statistically significant main effects were followed up with post-hoc comparisons (Fisher’s LSD). We also calculated Cohen’s d_s_ of all possible pairwise group comparisons as a measure of effect size. For this we used the ESCI intro software^203^.

The same LMM was used to compare the EMG envelopes across groups and conditions. To this end, the EMG amplitude was averaged over two time windows within the gait cycle in each group: 1. during early swing between −0.4 to −0.25 s relative to heel strike (foot lift phase), 2. between 0.04 and 0.08 s (foot drop) relative to heel strike. Post-hoc comparisons (Fisher’s LSD) for significant main effects were also completed.

Subsequently, EMG amplitude during both time intervals was correlated with corticomuscular coherence over significant time-frequency windows within each of the groups (young, old and PD) and conditions (treadmill and overground).

## Acknowledgements

We thank Bridie O’Connell and Kathryn McIntosh for support with data collection.

