## Supplementary material for "Corticomuscular control of walking in older people and people with Parkinson’s disease"

**Table S1** Main effect of condition and mean estimates of temporal gait parameters

| Gait parameter | Overground | | Treadmill | | Condition effect | |
|---|---|---|---|---|---|---|
| | mean | SEM | mean | SEM | F | P |
| Walking speed (km/h) | 4.05 | 0.065 | 4.04 | 0.065 | 1.03 | 0.31 |
| Stride time (s) | **1.104** | **0.01** | **1.067** | **0.01** | **30.84** | **<0.001** |
| Stride time variability (s) | 0.021 | 0.001 | 0.019 | 0.001 | 1.93 | 0.17 |
| Step time (s) | **0.545** | **0.005** | **0.529** | **0.005** | **22.93** | **<0.001** |
| Step time variability (s) | 0.011 | 0.001 | 0.011 | 0.001 | 0.002 | 0.96 |
| Stance phase (s) | **0.697** | **0.008** | **0.676** | **0.008** | **15.84** | **<0.001** |
| Swing phase/ Single support (s) | **0.395** | **0.004** | **0.384** | **0.004** | **6.93** | **0.01** |
| Double support (s) | 0.150 | 0.007 | 0.158 | 0.007 | 0.61 | 0.44 |

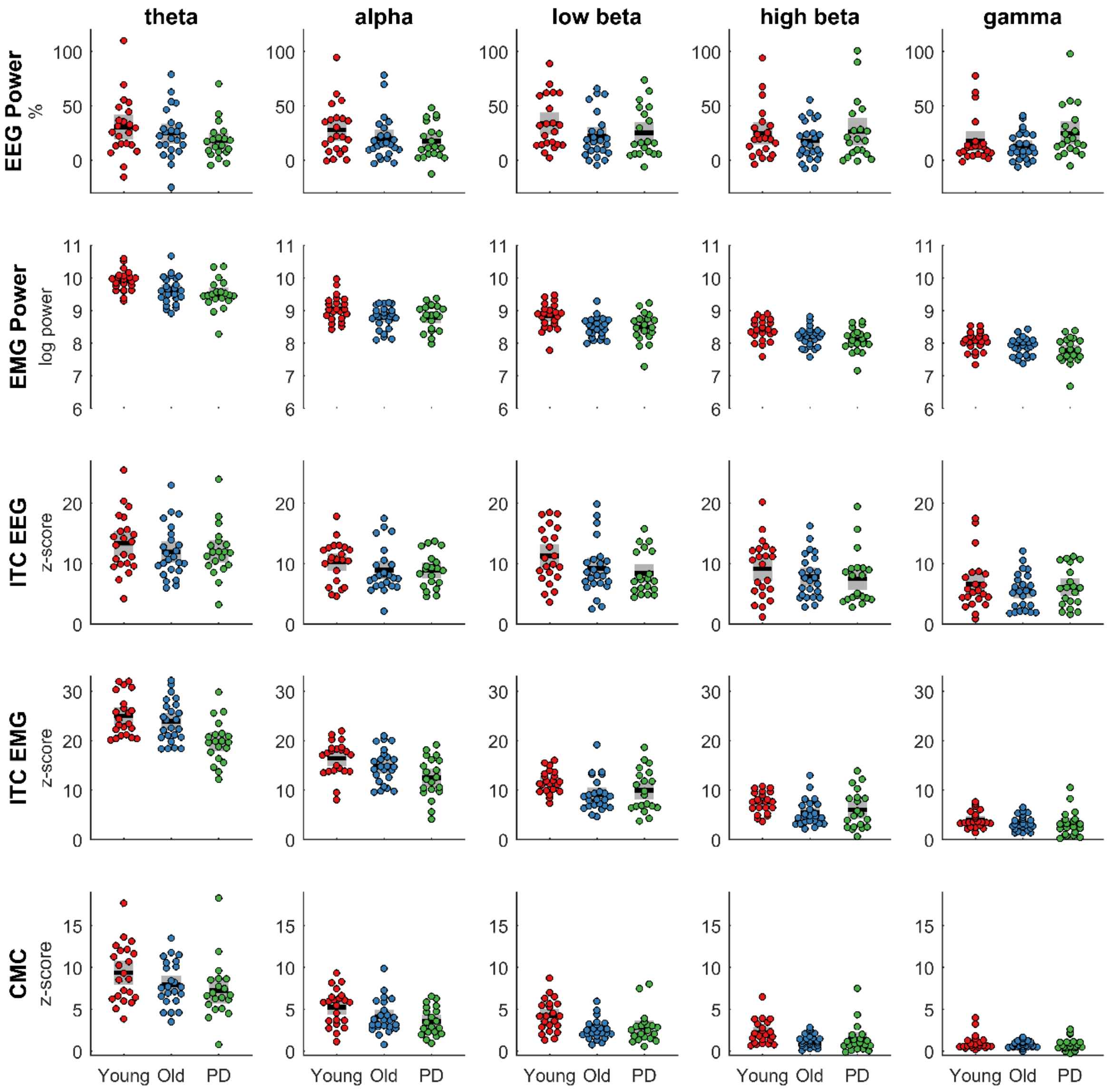


**Figure S2. Data of group comparison for spectral measures during overground walking.** Spectral measures (rows: EEG power, EMG power, ITC EEG, ITC EMG, CMC) were averaged over double support in different frequency bands (columns: theta, alpha, low beta, high beta, gamma) for each group (young, old, PD). Coloured dots show individual data of each participant, black horizontal lines show the group mean, grey boxes show the SEM. CMC, corticomuscular coherence; ITC, inter-trial coherence.

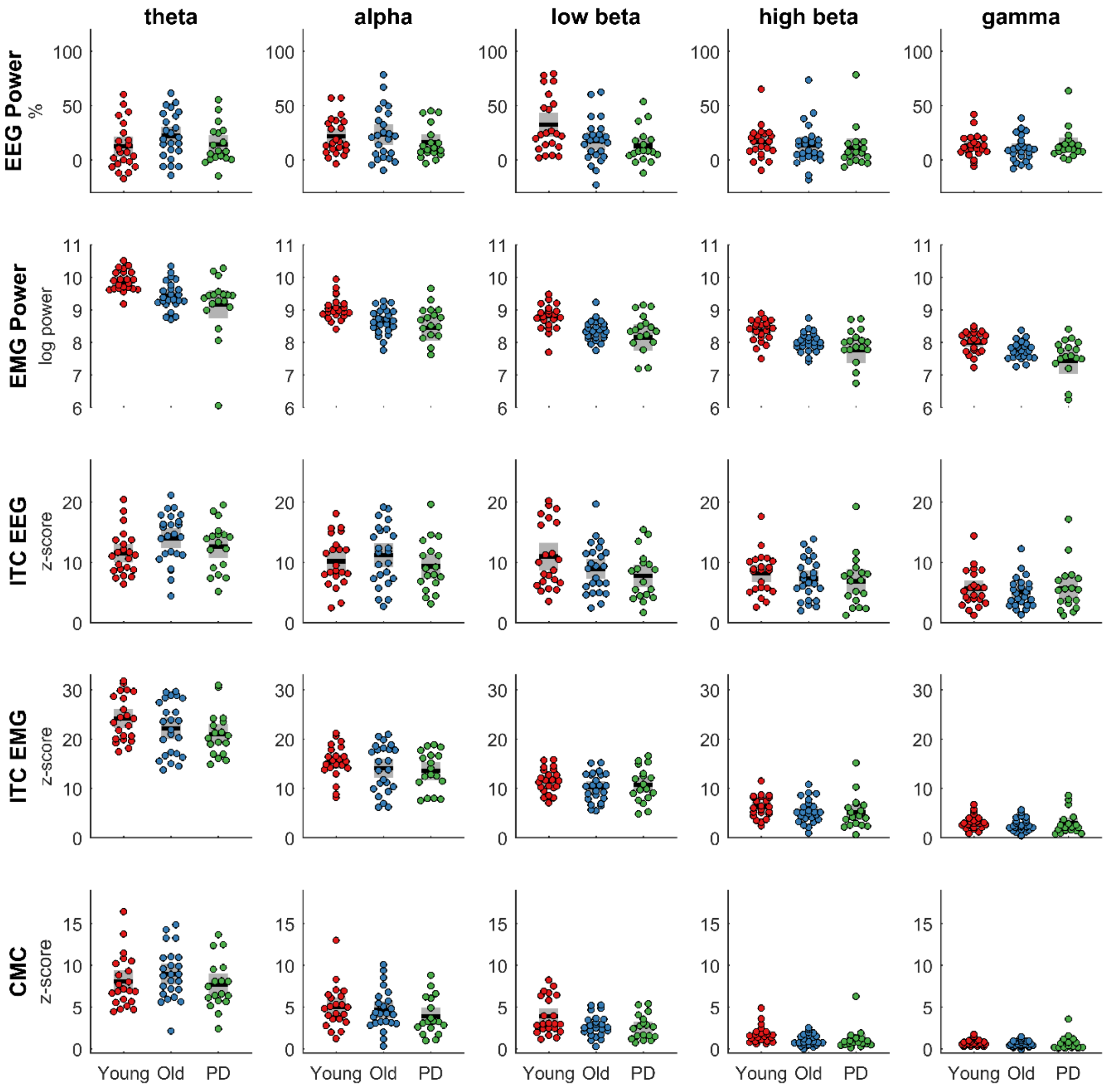


**Figure S3. Data of group comparison for spectral measures during treadmill walking.** Spectral measures (rows: EEG power, EMG power, ITC EEG, ITC EMG, CMC) were averaged over double support in different frequency bands (columns: theta, alpha, low beta, high beta, gamma) for each group (young, old, PD). Coloured dots show individual data of each participant, black horizontal lines show the group mean, grey boxes show the SEM. CMC, corticomuscular coherence; ITC, inter-trial coherence.

**Table S4** Main effect of condition for spectral measures and mean estimates with 95% confidence intervals

| Spectral measure | Overground | | Treadmill | | Condition effect | | |
|---|---|---|---|---|---|---|---|
| | mean | 95% CI | mean | 95% CI | F | P | $P_{adjusted}$ |
| **EEG power (%)** | | | | | | | |
| Theta | 24.5 | 19.1-29.8 | 16.4 | 11.0-21.9 | **5.53** | **0.020** | **0.042** |
| Alpha | 21.8 | 16.8-26.8 | 21.1 | 16.0-26.2 | 0.07 | 0.79 | 0.82 |
| Low beta | 27.1 | 21.6-32.6 | 21.5 | 16.0-27.1 | 3.38 | 0.068 | 0.13 |
| High beta | 22.9 | 17.9-27.9 | 14.0 | 8.9-19.1 | **11.1** | **0.001** | **0.004** |
| Gamma | 18.7 | 14.8-22.5 | 12.8 | 8.9-16.7 | **6.81** | **0.010** | **0.025** |
| **EMG power (log)** | | | | | | | |
| Theta | 9.7 | 9.6-9.8 | 9.5 | 9.4-9.6 | **11.5** | **0.001** | **0.004** |
| Alpha | 8.9 | 8.8-9.0 | 8.7 | 8.6-8.9 | **11.5** | **0.001** | **0.004** |
| Low beta | 8.6 | 8.5-8.7 | 8.4 | 8.3-8.6 | **13.8** | **<0.0005** | **0.004** |
| High beta | 8.2 | 8.1-8.3 | 8.1 | 8.0-8.2 | **19.8** | **<0.0005** | **0.004** |
| Gamma | 7.9 | 7.8-8.0 | 7.7 | 7.6-7.8 | **17.7** | **<0.0005** | **0.004** |
| **ITC EEG (z-score)** | | | | | | | |
| Theta | 12.4 | 11.4-13.4 | 12.7 | 11.7-13.8 | 0.25 | 0.62 | 0.70 |
| Alpha | 9.4 | 8.4-10.3 | 10.3 | 9.4-11.3 | 2.99 | 0.085 | 0.15 |
| Low beta | 9.6 | 8.6-10.7 | 9.3 | 8.2-10.3 | 0.36 | 0.55 | 0.65 |
| High beta | 8.2 | 7.2-9.1 | 7.4 | 6.5-8.4 | 2.53 | 0.11 | 0.18 |
| Gamma | 6.0 | 5.3-6.8 | 5.4 | 4.6-6.2 | 2.30 | 0.13 | 0.20 |
| **ITC EMG (z-score)** | | | | | | | |
| Theta | 22.9 | 21.8-24.1 | 22.4 | 21.2-23.5 | 1.55 | 0.22 | 0.31 |
| Alpha | 14.6 | 13.7-15.6 | 14.3 | 13.4-15.3 | 0.43 | 0.51 | 0.64 |
| Low beta | 10.3 | 9.5-11.0 | 10.8 | 10.1-11.6 | 2.18 | 0.14 | 0.21 |
| High beta | 6.3 | 5.6-7.0 | 5.6 | 4.9-6.3 | **7.02** | **0.009** | **0.025** |
| Gamma | 3.5 | 3.1-4.0 | 2.9 | 2.4-3.4 | **10.5** | **0.001** | **0.004** |
| **CMC (z-score)** | | | | | | | |
| Theta | 8.2 | 7.4-8.9 | 8.2 | 7.5-9.0 | 0.02 | 0.89 | 0.89 |
| Alpha | 4.3 | 3.8-4.9 | 4.6 | 4.0-5.1 | 0.64 | 0.42 | 0.55 |
| Low beta | 3.2 | 2.8-3.6 | 3.1 | 2.7-3.5 | 0.18 | 0.67 | 0.73 |
| High beta | 1.7 | 1.4-1.9 | 1.3 | 1.0-1.6 | **6.15** | **0.014** | **0.032** |
| Gamma | 0.9 | 0.7-1.0 | 0.7 | 0.5-0.8 | **6.92** | **0.009** | **0.025** |

CI, confidence interval; ITC, inter-trial coherence; CMC, corticomuscular coherence.

**Table S5** Post-hoc comparisons of significant group effects

| Spectral measure | Healthy young | | Healthy old | | PD | | P-value pairwise comparisons | | |
|---|---|---|---|---|---|---|---|---|---|
| | mean | 95% CI | mean | 95% CI | mean | 95% CI | Young-old | Young-PD | Old-PD |
| **EMG power (log)** | | | | | | | | | |
| Theta | 9.9 | 9.7-10.1 | 9.5 | 9.3-9.7 | 9.3 | 9.1-9.5 | **0.007** | **0.0002** | 0.18 |
| Alpha | 9.1 | 8.9-9.2 | 8.7 | 8.5-8.9 | 8.6 | 8.4-8.8 | **0.01** | **0.002** | 0.5 |
| Low beta | 8.8 | 8.6-9.0 | 8.4 | 8.3-8.6 | 8.3 | 8.1-8.5 | **0.004** | **0.001** | 0.46 |
| High beta | 8.4 | 8.2-8.5 | 8.1 | 7.9-8.3 | 7.9 | 7.8-8.1 | **0.02** | **0.001** | 0.19 |
| Gamma | 8.0 | 7.9-8.2 | 7.8 | 7.7-8.0 | 7.6 | 7.4-7.8 | 0.07 | **0.001** | 0.08 |
| **ITC EMG (z-score)** | | | | | | | | | |
| Theta | 24.6 | 22.8-26.4 | 23.0 | 21.3-24.8 | 20.3 | 18.4-22.2 | 0.2 | **0.002** | **0.04** |
| **CMC (z-score)** | | | | | | | | | |
| Low beta | 4.1 | 3.5-4.7 | 2.7 | 2.1-3.2 | 2.7 | 2.1-3.3 | **0.001** | **0.002** | 0.9 |

CI, confidence interval; ITC, inter-trial coherence; CMC, corticomuscular coherence.

**Table S6** Effect sizes (Cohen's $d_s$) of pairwise group comparisons

| Spectral measure | Young-Old | | Young-PD | | Old-PD | |
|---|---|---|---|---|---|---|
| | $d_s$ | 95% CI | $d_s$ | 95% CI | $d_s$ | 95% CI |
| **EEG power (%)** | | | | | | |
| Theta | 0.11 | -0.47, 0.69 | -0.33 | -0.95, 0.3 | -0.44 | -1.06, 0.18 |
| Alpha | -0.18 | -0.76, 0.40 | -0.42 | -1.05, 0.21 | -0.24 | -0.85, 0.38 |
| Low beta | -0.70 | -1.30, -0.10 | -0.74 | -1.37, -0.09 | -0.02 | -0.63, 0.59 |
| High beta | -0.28 | -0.90, 0.31 | -0.10 | -0.72, 0.53 | 0.18 | -0.43, 0.79 |
| Gamma | -0.29 | -0.90, 0.30 | 0.34 | -0.27, 0.97 | 0.64 | 0.00, 1.26 |
| **EMG power (log)** | | | | | | |
| Theta | -0.82 | -1.42, -0.21, | -1.22 | -1.89, -0.53 | -0.41 | -1.02, 0.21 |
| Alpha | -0.79 | -1.38, -0.18 | -0.99 | -1.60, -0.32 | -0.20 | -0.81, 0.41 |
| Low beta | -0.89 | -1.49, -0.28 | -1.11 | -1.78, -0.43 | -0.22 | -0.84, 0.39 |
| High beta | -0.71 | -1.31, -0.11 | -1.12 | -1.78, -0.44 | -0.40 | -1.02, 0.22 |
| Gamma | -0.54 | -1.13, 0.05 | -1.09 | -1.75, -0.41 | -0.55 | -1.17, 0.08 |
| **ITC EEG (z-score)** | | | | | | |
| Theta | 0.16 | -0.42, 0.74 | -0.05 | -0.68, 0.57 | -0.22 | -0.83, 0.40 |
| Alpha | -0.05 | -0.626, 0.53 | -0.33 | -0.96, 0.30 | -0.29 | -0.90, 0.33 |
| Low beta | -0.60 | -1.19, 0.00 | -0.89 | -1.54, -0.23 | -0.28 | -0.89, 0.33 |
| High beta | -0.32 | -0.90, 0.26 | -0.45 | -1.08, 0.19 | -0.12 | -0.73, 0.49 |
| Gamma | -0.33 | -0.91, 0.26 | -0.10 | -0.72, 0.53 | 0.24 | -0.38, 0.85 |
| **ITC EMG (z-score)** | | | | | | |
| Theta | -0.37 | -0.95, 0.22 | -1.00 | -1.66, -0.34 | -0.64 | -1.26, -0.01 |
| Alpha | -0.42 | -1.01, 0.17 | -0.85 | -1.50, -0.19 | -0.42 | -1.03, 0.20 |
| Low beta | -0.71 | -1.30, -0.11 | -0.50 | -1.13, 0.14 | 0.22 | -0.39, 0.83 |
| High beta | -0.58 | -1.17, 0.01 | -0.54 | -1.17, 0.10 | 0.06 | -0.55, 0.67 |
| Gamma | -0.38 | -0.96, 0.21 | -0.31 | -0.94, 0.32 | 0.07 | -0.54, 0.68 |
| **CMC (z-score)** | | | | | | |
| Theta | -0.11 | -0.69, 0.47 | -0.53 | -1.16, 0.11 | -0.41 | -1.03, 0.21 |
| Alpha | -0.40 | -0.98, 0.19 | -0.78 | -1.42, -0.13 | -0.37 | -0.99, 0.25 |
| Low beta | -1.05 | -1.67, -0.43 | -1.01 | -1.66, -0.34 | -0.04 | -0.65, 0.57 |
| High beta | -0.78 | -1.37, -0.17 | -0.67 | -1.30, -0.02 | 0.13 | -0.49, 0.74 |
| Gamma | -0.38 | -0.96, 0.21 | -0.28 | -0.90, 0.35 | 0.11 | -0.50, 0.72 |

CI, confidence interval; ITC, inter-trial coherence; CMC, corticomuscular coherence.

**Table S7** Main effect of condition for EMG envelopes and mean estimates with 95% confidence intervals

| **EMG envelope amplitude (%) at gait cycle phase** | **Overground** | | **Treadmill** | | **Condition effect** | |
|---|---|---|---|---|---|---|
| | mean | 95% CI | mean | 95% CI | F | P |
| Foot drop | 57.1 | 52.7, 61.4 | 51.9 | 47.6, 56.3 | **9.26** | **0.003** |
| Foot lift | 58.1 | 55.1, 61.1 | 59.8 | 56.8, 62.9 | 2.42 | 0.12 |

CI, confidence interval.

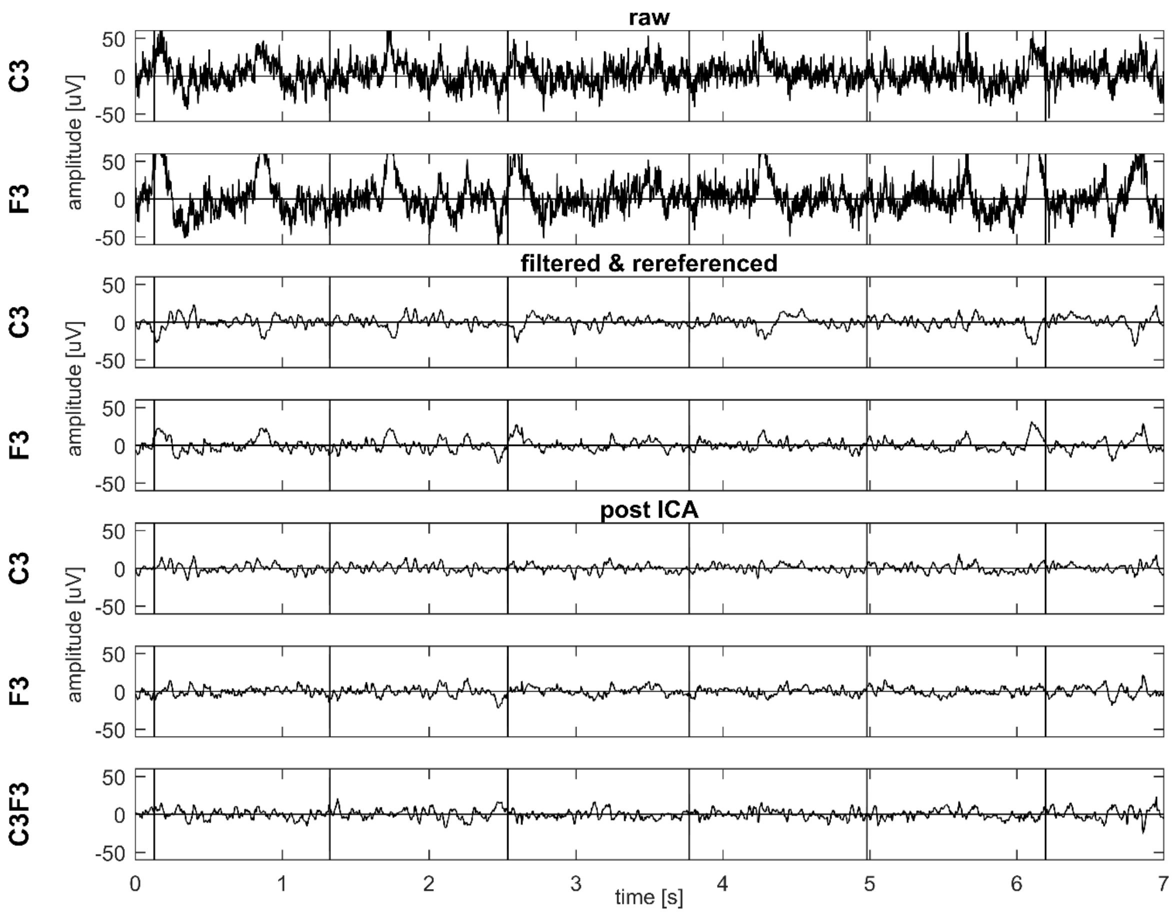


**Figure S8. Data of one healthy older participant during overground walking.** EEG signals of the C3 and F3 channels at different pre-processing steps: raw C3 and F3 signals (two top panels), after filtering and re-referencing (third and fourth panel), after Independent Component Analysis (ICA; fifth and sixth panel), bipolar signal C3-F3 (bottom panel). The vertical lines present heel strikes of the right foot.
